# *GeneSwitches* : Ordering gene-expression and functional events in single-cell experiments

**DOI:** 10.1101/832626

**Authors:** Elaine Y. Cao, John F. Ouyang, Owen J.L. Rackham

## Abstract

**Summary:** Emerging single-cell RNA-seq technologies has made it possible to capture and assess the gene expression of individual cells. Based on the similarity of gene expression profiles, many tools have been developed to generate an in silico ordering of cells in the form of pseudo-time trajectories. However, these tools do not provide a means to find the ordering of critical gene expression changes over pseudo-time. We present GeneSwitches, a tool that takes any single-cell pseudo-time trajectory and determines the precise order of gene-expression and functional-event changes over time. GeneSwitches uses a statistical framework based on logistic regression to identify the order in which genes are either switched on or off along pseudo-time. With this information, users can identify the order in which surface markers appear, investigate how functional ontologies are gained or lost over time, and compare the ordering of switching genes from two related pseudo-temporal processes.

**Availability:** GeneSwitches is available at https://geneswitches.ddnetbio.com

**Supplementary Information:** is available at http://www.ddnetbio.com/files/GeneSwitches_SI.pdf

## 1 Introduction

Since the advent of next-generation sequencing, one major application has been the study of molecular changes that take place during cellular transitions, such as those that follow an external stimulus or during a cell conversion. Typically bulk RNA sequencing data (RNA-seq) would be produced at various time points after stimulation in order to capture gene expression changes that underpin the transition. However, in order to accurately capture the time-dependent gene expression changes, it is necessary to sample RNA-seq at a high resolution relative to the time-scale of the transition. This can be both expensive and difficult to implement. Furthermore, because RNA-seq samples gene expression from a population of cells, the accuracy of this technique often relies greatly on the progression of the transition being relatively stable across the population, particularly where no surface markers for intermediate states are known which can be used to FACs purify the populations prior to sequencing. As a result, these experiments tend to not capture the true time resolution of the process and can be confounded by bifurcating or mixed populations of cells.

In recent years the introduction of single-cell RNA-seq (scRNA-seq) has provided a means to circumvent these issues. With the ability to capture and assess the level of gene expression in individual cells comes the ability to order cells over pseudo-time. Pseudo-time provides an *in silico* ordering of cells (or trajectory) based on a comparison of their gene expression profiles. Many tools have been developed for this, for instance, Monocle and Slingshot (reviewed in Tian et al., 2019). However, extracting the underlying order of gene expression changes from these trajectories can be difficult. However, being able to interpret these gene expression changes in terms of the order that they occur would allow for a fuller understanding of the underlying biological processes. To address this, here we developed GeneSwitches, a statistical framework that processes scRNA-seq data together with a pseudo-time trajectory to find the set of genes that switch during the transition. Furthermore, for each gene, we calculate a switching time and associated confidence level. With this information, it is possible to (1) investigate how gene-regulatory networks or gene ontologies are gained or lost over time, (2) stratify selected gene sets (e.g. surface markers) by the order in which they appear and (3) identify key differences in the gene-expression changes in cell transitions that bifurcate over time.

To demonstrate this we apply GeneSwitches to scRNA-seq data from the differentiation of human embryonic stem cells (hESC) to cardiomyocytes (CM) (Friedman *et al*., 2018), highlighting both ontological and phenotypical ordering of events (Fig. 1).

**Fig. 1.**
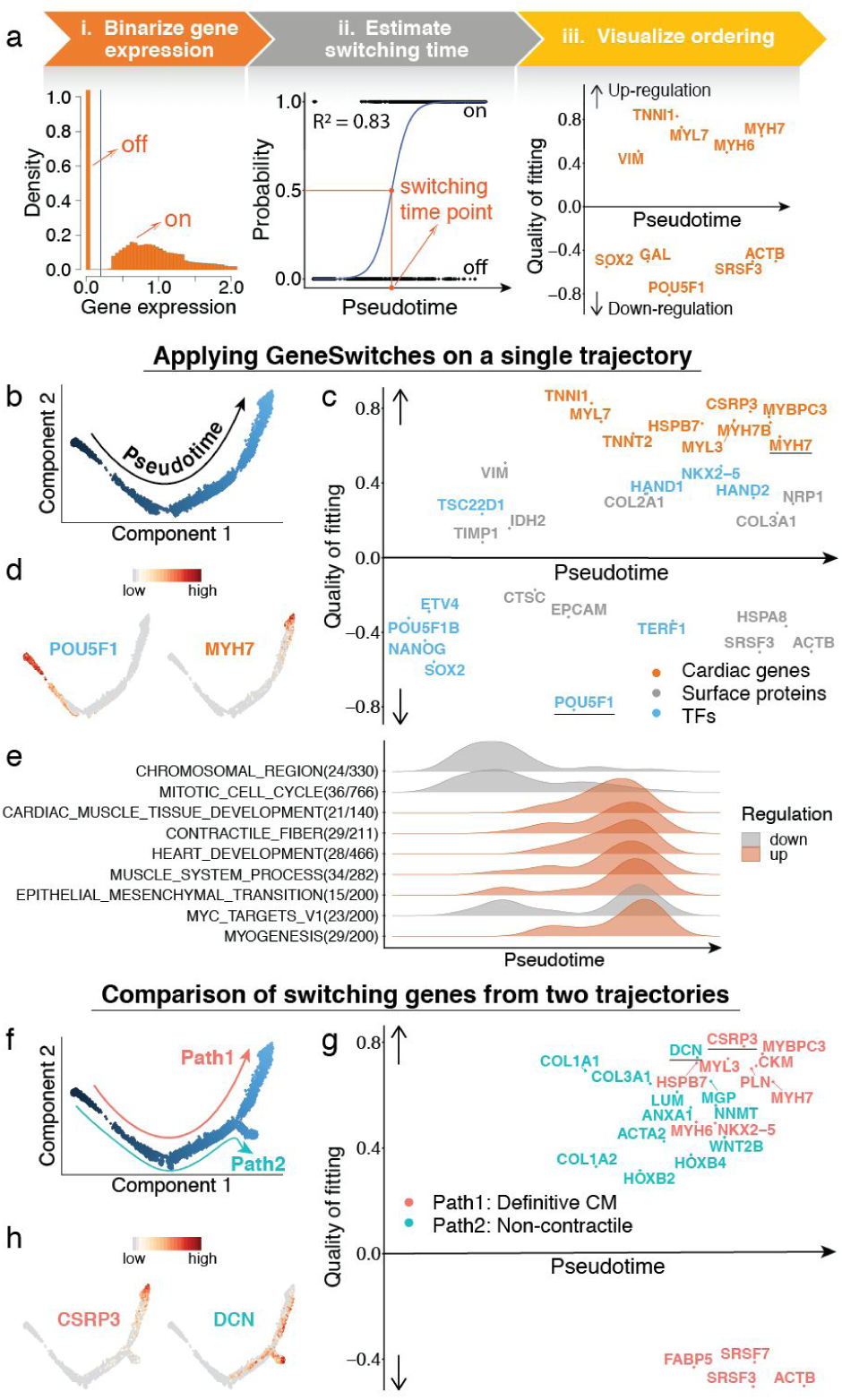
GeneSwitches overview and example analysis. **a**) Key steps of GeneSwitches **i**) Histogram of expression of all the genes in all cells and the global threshold for the on/off state (blue line). **ii**) Logistic regression is being fitted to the binarized data for each gene and the switching time is defined where the probability is 0.5. The quality of fit is defined by McFadden’s Pseudo R^2^. **iii**) Visualisation of switching genes along the pseudo-time. **b**) Linear trajectory inferred by Monocle2 for the differentiation of hESC to CM. **c**) Visualisation of the order of the top switching genes from various sets of known proteins. **d**) Expression of example genes from (c). **e**) Density plots of switching genes for significantly over-represented functional ontologies. **f**) The inferred trajectory shows a bifurcation of cell fates. **g**) Visualisation of distinct switching genes from the two paths filtered by the McFadden’s Pseudo R^2^. **h**) Expression of example genes from (g).

## 2 GeneSwitches functions and examples

The main workflow of GeneSwitches and the biological insights that can be derived are summarised as follows:

### 2.1 Data preprocessing

GeneSwitches requires two inputs, namely the gene expression matrix and the pseudo-time ordering of each single cell. First, GeneSwitches binarizes the gene expression into either an “on” or “off” state to enable the identification of switching events. To do this we plot the expression distribution of all the genes in all cells and look for a separation between the zero and expressed distributions to identify a global threshold separating the “on” and “off” gene expression states (Fig. 1a,i). Once identified this threshold is applied to the gene expression matrix to generate a binary gene expression matrix.

### 2.2 Ordering and visualisation of switching genes

Next, the binarized gene expression is used as a dependent variable in logistic regression with the pseudo-time value of each cell providing the independent variable. In doing so the probability of expression throughout pseudo-time is calculated and the quality of fit is determined by McFadden’s Pseudo R^2^. Following this, a set of switching genes is extracted (see supplementary methods) and for each the pseudo-time where the probability is 0.5 is defined as their switching time (Fig. 1a,ii). These switching genes are then visualised in order to describe the pseudo-time process in terms of the gene-expression events (Fig.1a,iii).

### 2.3 Ordering and visualisation of gene classes and functional groups

Switching genes can be used to investigate the functional nature of the pseudo-time trajectory. For instance, it might be desirable to know for a set of known surface proteins at what point they are activated or deactivated during a transition in order to facilitate the identification of suitable markers on which to sort cells that are transitioning. GeneSwitches can also identify the order in which functional ontologies are acquired or lost during a transition. To visualise these changes we provide the functionality to plot the density of switching genes from each ontological class with respect to pseudo-time in order to study when and how different functional classes are important.

As an example, we calculated the pseudo-time trajectory for the differentiation of hESC to cardiomyocytes (CM) using Monocle2 (Trapnell et al., 2014; Qiu et al., 2017) (Fig. 1b). Following this, GeneSwitches identified that TIMP1 and VIM were early surface proteins to be activated, indicating that they might represent good candidate markers to identify cells progressing along the differentiation process more rapidly (Fig. 1c). Furthermore, we also observe that POU5F1 is deactivated early, whilst MYH7 is activated late (Fig. 1d). Functional ontology analysis showed that the cell-cycle related ontologies were down-regulated at an early time and cells acquired cardiac-related functions later in the pseudo-time (Fig.1e).

### 2.4 Comparing the ordering from two related pseudo-time processes

If GeneSwitches has been used to analyse two related pseudo-time trajectories it is possible to compare the switching genes and their switching times. For instance, it is often the case in differentiation processes that certain populations of cells bifurcate, with each group being committed to a different cell state. GeneSwitches can be used to compare these trajectories, looking for similarities and differences in the switching genes, as well as their switching times. Once identified these similarities and differences can be used to better understand the molecular events that predispose each population to their fate choice. For instance, cells undergoing differentiation from hESCs toward a cardiomyocyte (CM) fate bifurcate, giving rise to one path progressing towards definitive CM state and a second path towards a non-contractile cardiac derivative state (Fig. 1f). GeneSwitches identified distinct switching events for the two paths (Fig. 1g), with the definitive CM path gaining cardiac markers, namely CSRP3 and NKX2-5, while the non-contractile path gaining DCN and COL1A2 (Fig. 1h).

In summary, GeneSwitches can help identify the timing of gene expression events within a pseudo-time trajectory, which in turn allows for a fuller understanding of the order of regulatory and functional events that occur during a cellular transition.

## Funding and Acknowledgments

O.J.L.R and J.F.O conceived the project, designed the study. and oversaw its delivery. E.Y.C developed the code, implemented the algorithm and executed the biological studies. All authors wrote the paper. This work has been supported by a Singapore National Research Foundation competitive research grant (NRF-CRP20-2017-0002).

