## Supplemental Methods for "*GeneSwitches* : Ordering gene-expression and functional events in single-cell experiments"

The goal of GeneSwitches is to discover the order of gene expression and functional events during cell state transitions at a single-cell resolution. It works on any single-cell trajectory or pseudo-time ordering of cells to discover the genes that act as on/off switches between cell states and importantly the ordering at which these switches take place.

#### S1. General workflow of GeneSwitches

##### S1.1. Input datasets

GeneSwitches requires two inputs, namely a gene expression matrix and corresponding pseudo-time ordering of each cell. Log-normalized gene expression is recommended. Dimensionality reduction (e.g. PCA or UMAP) input is optional and only needed to visualise gene expression changes. Apart from the direct input of this information, GeneSwitches also provides functions to convert trajectory inference results from published methods, namely Monocle2 (Trapnell et al., 2014; Qiu et al., 2017) and Slingshot (Street et al., 2018). In order to do this, the states of the desired path are required to convert Monocle 2 objects, while the name of the desired pseudo-time path is required for Slingshot.

##### S1.2. Binarize gene expression

GeneSwitches first binarizes the input gene expression into either an "on" or "off" gene-expression state to enable the identification of switching events. To achieve this, the expression distribution of all the genes in all cells is plotted in order to identify a separation between the zero and expressed distributions which is then used as a global threshold between states (Fig. S1; 0.2 is the default value for log-transformed gene expression data). This threshold is then applied to the gene expression matrix to generate a binary state matrix.

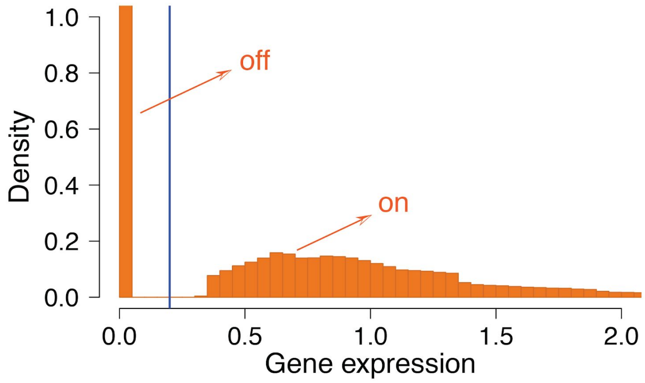


***Fig S1.*** *Histogram of log-transformed gene expression of all the genes in all cells. The blue line indicates the global threshold for the on/off state (0.2 is the default value).*

#### S1.3. Fit logistic regression & estimate switching time

Logistic regression is a statistical model that is used to explain the relationship between one dependent binary variable and other variables. In order to accurately fit a logistic model the availability of a high number of data points is required, something that is easily achievable with scRNA-seq data. Accordingly, for each gene in each cell, the binarized state is used as a dependent variable in logistic regression with the pseudo-time value providing the independent variable. Using this, GeneSwitches calculates the probability of gene-expression throughout pseudo-time.

GeneSwitches uses the fastGLM (Marschner, 2011) package in R to fit the logistic model. The quality of fit is estimated using McFadden's Pseudo $R^2$ (McFadden, 1972) and the Benjamini-Hochberg procedure is performed across all genes to calculate the FDR. Following this, the switching pseudo-time is determined by the point at which the fitted model crosses a probability threshold of 0.5 (Fig. S2). For genes with a high dropout rate (i.e. an observed gene expression of zero), GeneSwitches allows the user to use a random downsampling of zero expression values before fitting in order to rescue the prediction of switching times (see section S6. for more details).

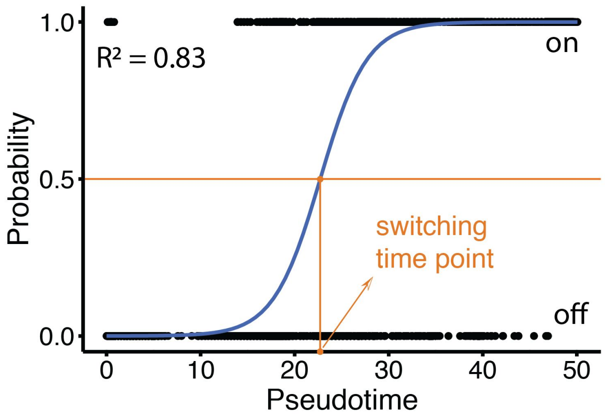


***Fig S2.*** *Logistic regression fitted to the binarized data for each gene: The switching time is determined by the point at which the fitted (blue) model crosses the probability threshold of 0.5. McFadden's Pseudo $R^2$ indicates the quality of fitting.*

#### S1.4. Visualize ordering of switching genes

A poor fit of the logistic regression to the observed data identifies those genes that don't display a switching behaviour over time and are often those genes that have constant expression, or a transitory expression pattern. In order to remove these genes and to determine the set of switching genes, GeneSwitches by default filters out the genes for which 1) the percentage of cells where the expression is 0 is greater than 90%, 2) the fitted logistic regression does not cross probability of 0.5, 3) the FDR is greater than 0.05 or 4) the pseudo $R^2$ is smaller than 0.03. The remaining genes are considered as switching genes and can be visualized along pseudo-time. Genes that are switched on are plotted above the pseudo-time line, while those switching off are below the line (Fig. S3).

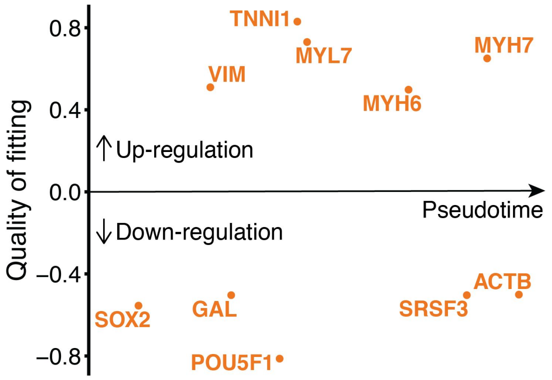


***Fig S3.*** *Visualisation of switching genes along the pseudo-time. The absolute value of the y-axis is the quality of fitting defined by McFadden's Pseudo $R^2$, and the positive and negative signs indicate up- and down-regulation respectively.*

One can also extract specific gene type(s) to visualise. By default, GeneSwitches provides gene type lists containing surface proteins (Bausch-Fluck et al., 2015) and transcription factors (Lambert et al., 2018), but users can pass their own gene type lists. This is done by providing a data frame, namely *my_genetypes,* with rows as genes (non-duplicated) and two columns with name *genenames* and *genetypes* as parameter *genelists* to the function *filter_switchgenes*.

### S2. Trajectory inference for the differentiation of human embryonic stem cells

To demonstrate the general workflow, we applied GeneSwitches to scRNA-seq data (39,794 cells in total) from the differentiation of human embryonic stem cells (hESC) to cardiomyocytes (CM) (Friedman et al., 2018). In order to do this, we first calculated the pseudo-time trajectory using Monocle2. It shows a bifurcating cell fate of cardiac hESC differentiation that gives rise to definitive cardiomyocytes (Path1) or non-contractile cardiac derivatives (Path2). Path1 consists of cells from states 1, 2, 4 and 5, while Path2 consists of those from states 1, 2, 4 and 6 (see Fig. S4). The function *convert_monocle2* can then be used with the identified Monocle states to create the GeneSwitches input objects for the two paths.

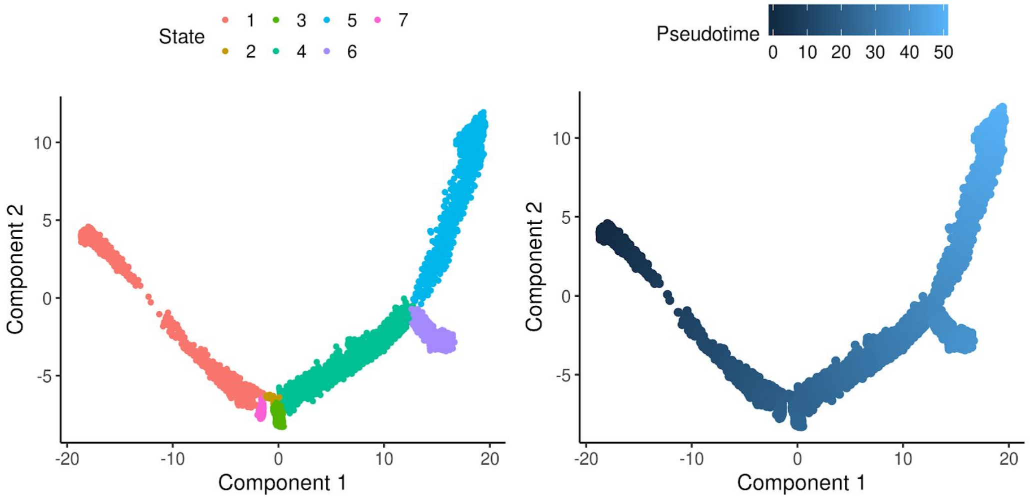

***Fig S4.*** *Trajectory inferred using Monocle2 for all 39,794 cells. The left panel is colored by states, which denote branches and can be used to extract paths. The right panel is colored by pseudo-time, showing that cells develop from state 1 to the bifurcation point that gives rise to states 5 and 6 respectively.*

### S3. GeneSwitches applied to scRNA from hESCs differentiated to definitive cardiomyocytes (Path1)

GeneSwitches was first applied to the single trajectory in which cells differentiate from hESC to definitive cardiomyocytes (Path1). There are in total of 28,852 cells in this path. The gene expression information from these cells was processed as outlined in section S1, the default global threshold 0.2 was used to binarize the gene expression and a logistic regression fitting with downsampling conducted. After identifying the switching genes, the overall top 20 best fitting switching genes (according to the pseudo $R^2$) and top 15 either surface proteins or TFs were extracted for visualization (Fig. S5).

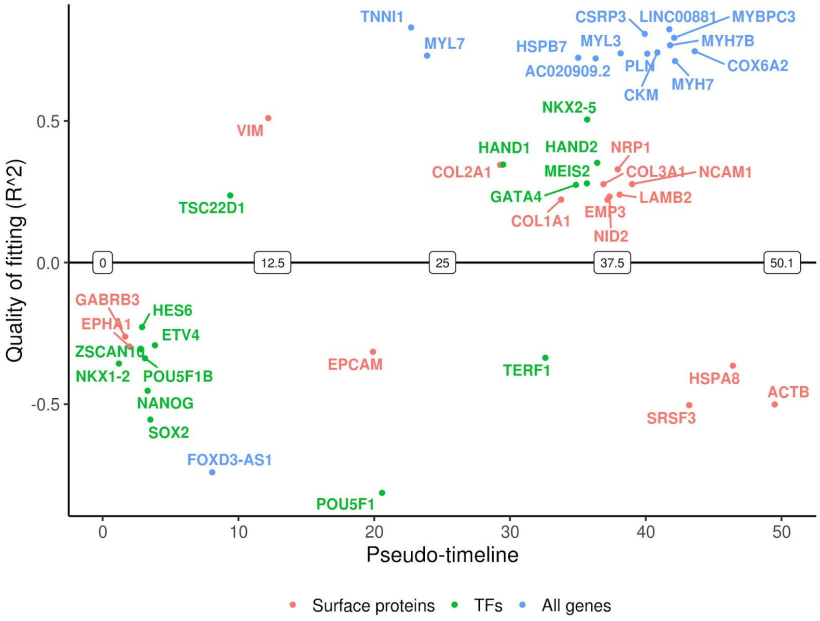


***Fig S5.*** *Visualisation of the order of the top best fitting switching genes as-well-as the top surface proteins or transcription factors (TFs).*

VIM was identified as a surface protein activated early, indicating that it might represent a good candidate marker for quantifying cells progressing along the differentiation process more rapidly. Furthermore, we also observe that POU5F1 is deactivated after SOX2 and NANOG indicating that the parts of the pluripotency network regulated by POU5F1 are the last to be switched off in this differentiation.

### S4. Order pathways along the pseudo-timeline

GeneSwitches can also identify the order in which functional ontologies or pathways are acquired or lost during a transition. By default, GeneSwitches includes the pathways provided by MSigDB hallmark

(Liberzon,A. et al., 2015), C2 KEGG (Kanehisa and Goto, 2000) and C5 gene ontology geneset collections. Alternatively, users can provide a list of functional ontologies that are more suitable for their system as parameter *pathways* to the function *find_switch_pathway*.

Switching genes from each pathway are identified and a hypergeometric test is applied to extract the pathways that are significantly overrepresented. The switching time of the pathway is then determined by calculating the median switching time of genes in that pathway. To better visualise the functional changes, ridge plots show the density of switching genes from the same pathway along the pseudo-time.

Functional ontology analysis of Path1 showed that the cell-cycle related ontologies were down-regulated at an early time and cells acquired cardiac-related functions later in the pseudo-time (Fig. S6).

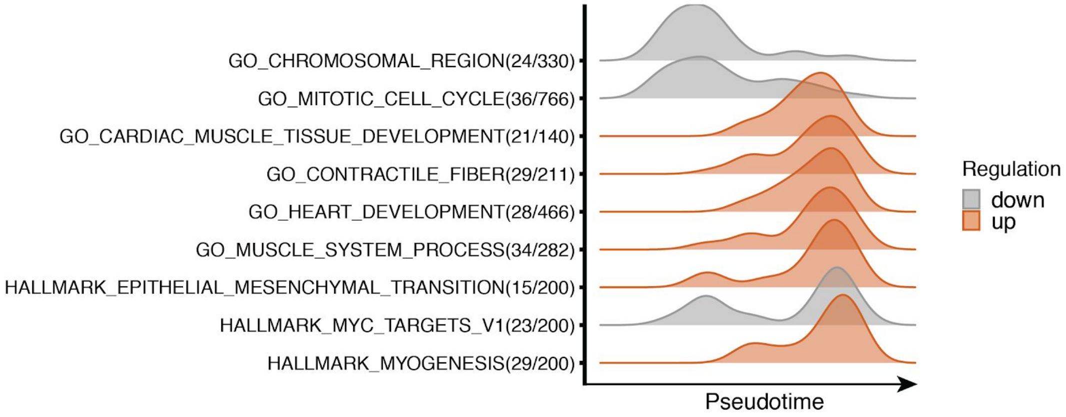


***Fig S6.*** *Density plots of switching genes for significantly over-represented functional ontologies.*

### S5. Comparing switching genes from two trajectories

If GeneSwitches has been used to analyse two related pseudo-time trajectories it is possible to compare the list of switching genes and their switching times. In order to do this, the same criteria should be used to identify significant switching genes for each of the two trajectories. Following this, the switching genes that are common between two trajectories can be plotted. More importantly, GeneSwitches can determine the switching genes that are specific to each trajectory and plot these on one timeline. GeneSwitches can also scale the timelines to be the same length (default number of bins is 100) so that the observed differences can be considered based on the percentage of the trajectory covered rather than actual pseudo-time.

To demonstrate this we compare the two paths observed in the cardiac hESC differentiation dataset described above. First, we processed the cells from Path2 (non-contractile cardiac derivatives) in the same ways as Path1 to identify the associated switching genes. GeneSwitches can then be used to find distinct switching events for these two paths (Fig. S7), with the definitive CM path gaining cardiac markers, such as CSRP3, while the non-contractile path gaining DCN. The identification of the differences along with their timings can facilitate follow up experiments to find the determinants of these bifurcations.

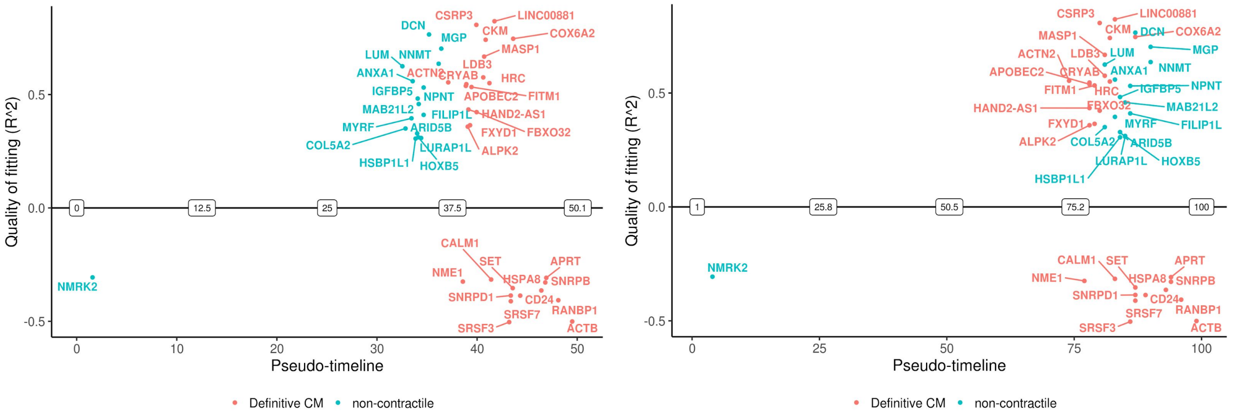


***Fig S7.*** *Visualisation of top 15 best fitting distinct switching genes from the two paths without (left) or with (right) timeline scaling.*

### S6. Downsampling of zero expression for the high-dropout cells

Owing to technical limitations and the low amount of starting material, scRNA-seq can be affected by the high number of dropouts. For example, gene GUCY1A1 was not detected (zero expression) in 84.7% of cells in Path1 and cells started to acquire this gene at a late stage. Fitting a logistic regression with data with high dropout results in a failure to identify the switching time because the large number of zeros deflates the probability estimation (Fig. S8a).

In this case, GeneSwitches can first apply a random downsampling of zero expression so that the remaining cells contained 50% of zero-expressed cells. Following this, by fitting logistic regression it is possible to identify the switching time of GUCY1A1 as being at a pseudo-time of 27 (Fig. S8b). Hence, random downsampling of zero expression before fitting the logistic model is able to recover the switching time of lowly expressed genes.

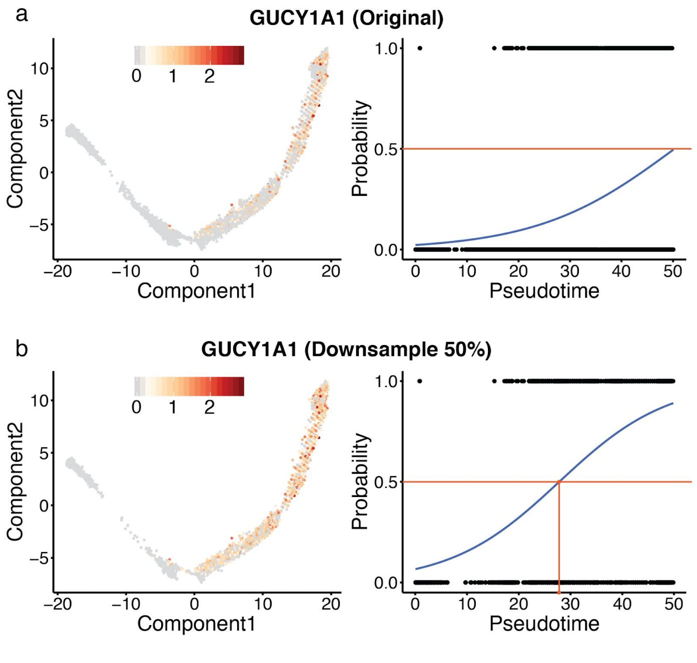

*Fig S8.* *Downsampling of zero expression for the high-dropout cells. a) Gene GUCY1A1 was not detected (zero expression) in 84.7% of cells in Path1, and fitting to the logistic model failed to identify the switching time since the fitted line did not cross probability 0.5. b) Random downsample of zero expression so that the remaining cells contained 50% of zero-expressed cells for GUCY1A1. Then fitting logistic regression to the downsampled data, the switching time of GUCY1A1 was identified at pseudo-time 27.*

### S7. Robust ordering of switching genes

In order to show that the ordering of switching genes is robust against the number of cells, we subsample the cells in Path1 (~ 28k cells in total) into 17k, 10k, 5k, 2.5k, 1k, 500, 250 and 100 cells to apply GeneSwitches respectively. The Jaccard index was calculated to compare the reported significant switching genes between the original and each subsample cases. The Jaccard index measures the similarity for the two sets of data, with a range from 0 (not overlapping) to 1 (identical), the higher the index, the more similar the two data sets. We observed that subsamples with over 2.5k cells, which is approximately 10% of the original cells, had a high Jaccard index close to 90% (Fig. S9a).

The order of the top 60 switching genes identified in the original dataset was highly conserved in the subsamples with over 2.5k cells (Fig. S9b). As a result, we can conclude that GeneSwitches should be able to provide a reliable ordering of genes even in cases where the number of cells is low.

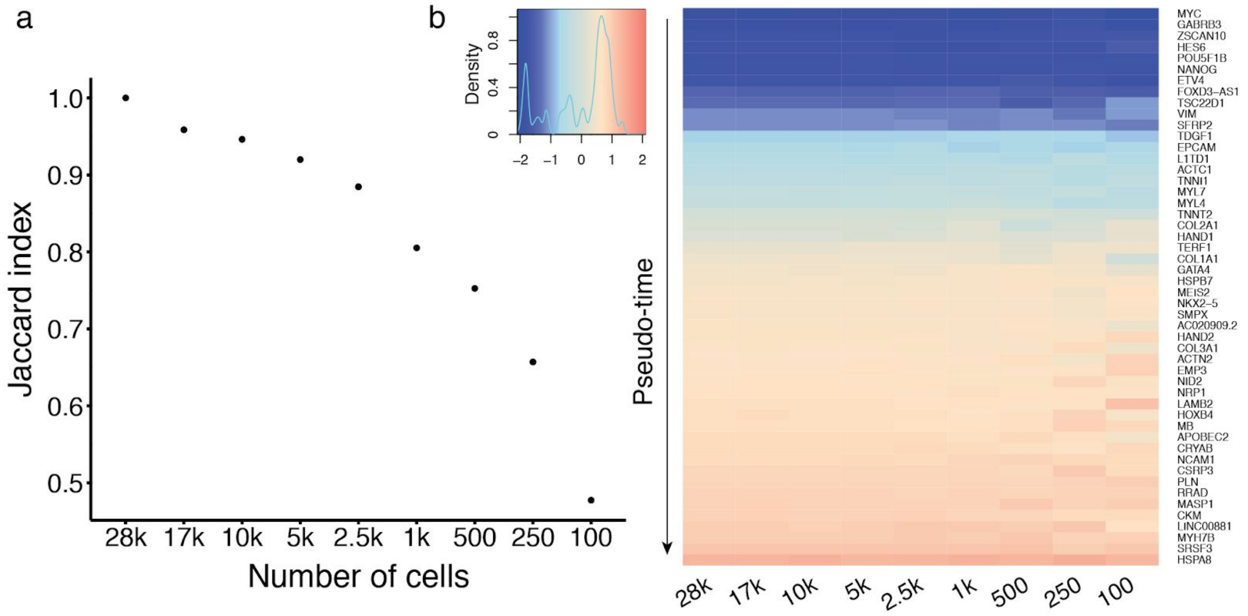


*Fig S9.* *Robust ordering of switching genes. a) The Jaccard index was calculated to compare the reported significant switching genes between the original and each subsample cases. We observed that subsamples with over 2.5k cells, which is approximately 10% of the original cells, had a high Jaccard index closed to 90%. b) Heatmap shows the order of the top 60 switching genes identified in the original dataset across each subsample case. The shades reflected that the ordering was highly conserved in the subsamples with over 2.5k cells.*
